# Divergence and introgression in small apes, the genus *Hylobates*, revealed by reduced representation sequencing

**DOI:** 10.1101/2020.05.31.126078

**Authors:** Kazunari Matsudaira, Takafumi Ishida

**Author notes:** Corresponding Author: Kazunari Matsudaira, Department of Biological Sciences, School of Science, The University of Tokyo, Hongo 7-3-1, Bunkyo-ku, Tokyo 113-0033, Japan.

## Abstract

Gibbons in the genus *Hylobates*, which live in Southeast Asia, show great diversity, comprising seven to nine species. Natural hybridisation has been observed in the species contact zones, although the roles played by hybridisation and introgression in the evolution of these species remain unclear. To uncover the divergence history and the contributions of hybridisation and introgression to the evolution of *Hylobates*, random amplicon sequencing-direct (GRAS-Di) analysis was employed to genotype 47 gibbons, representing eight species from three genera. After quality filtering, over 300,000 autosomal single-nucleotide variant (SNV) sites were identified. The SNV-based autosomal phylogeny, together with the mitochondrial phylogeny, supported a divergence pattern beginning approximately 4.3 million years ago. First, the mainland species, *H. pileatus* and *H. lar*, consecutively diverged from the Sundaic island species. Second, *H. moloch*, in Java (and likely *H. klossii*, in the Mentawai Islands) diverged from the other species. Third, *H. muelleri*, in Borneo, and *H. agilis*/*H. albibarbis*, in Sumatra and southwestern Borneo, diverged. Lastly, *H. agilis* and *H. albibarbis* diverged from each other. The Patterson’s D-statistics indicated significant introgression between *H. lar* and *H. pileatus*, between *H. lar* and *H. agilis*, and between *H. albibarbis* and *H. muelleri*, and weak introgression was identified between *H. moloch* and *H. albibarbis*, and between *H. moloch* and *H. muelleri abbotti*, suggesting incomplete reproductive barriers among *Hylobates* species and that hybridisation and introgression occur whenever the distribution ranges contact. Some candidates for introgressed genomic regions were detected, and the functions of these would be revealed by further genome-wide studies.

## Introduction

Primates is a highly diversified mammalian order, with an estimated 500 (and a minimum of 350) extant species (Groves 2001; Rylands and Mittermeier 2014). Recently, genome-wide studies of non-human primates have become possible, revealing evolutionary processes, such as speciation, and the demographic histories of various taxa (Prado-Martinez et al. 2013; Xue et al. 2016; Nater et al. 2017). Historical hybridisation and introgression have been detected in many cases (e.g., chimpanzees and bonobos [de Manuel et al. 2016], macaques [Fan et al. 2018], baboons [Rogers et al. 2019], and Dryas and green monkeys [van der Valk et al. 2020]); thus, these processes are recognised as common phenomena.

Moreover, the potential roles played by hybridisation and introgression in the evolution of non-human primates, such as adaptive introgression, have been widely discussed (Nye et al. 2018; Rogers et al. 2019; van der Valk et al. 2020), although further studies remain necessary to reveal how hybridisation occurs and the degree to which introgression has affected primate evolution. Among primates, Hylobatidae, which is a family of small apes or gibbons, is considered to be a good model for studying the contributions of divergence and hybridisation/introgression to evolution, especially as an analogy for evolution in hominins (Zichello 2018).

Gibbons are small apes that live in the Indomalaya realm and show great diversity among four extant genera, which each have different numbers of chromosomes (*Hoolock* [2n = 38], *Hylobates* [2n = 44], *Symphalangus* [2n = 50], and *Nomascus* [2n = 52]), comprising 15 to 20 species (Roos 2016; Fan et al. 2017). Among these four genera, *Hylobates* shows the highest taxonomic diversity, followed by *Nomascus*. *Hylobates* is distributed throughout both the mainland and islands of Southeast Asia, and seven allopatric species are recognised, and characterised by various pelage colours and vocalisations: *Hylobates pileatus* (pileated gibbons), *H. lar* (white-handed gibbons), *H. agilis* (agile gibbons), *H. albibarbis* (Bornean white-bearded gibbons), *H. muelleri* (Müller’s gibbons), *H. moloch* (Javan gibbons), and *H. klossii* (Kloss’s gibbons) (Fig. 1; Groves 2001).

**Fig. 1.**
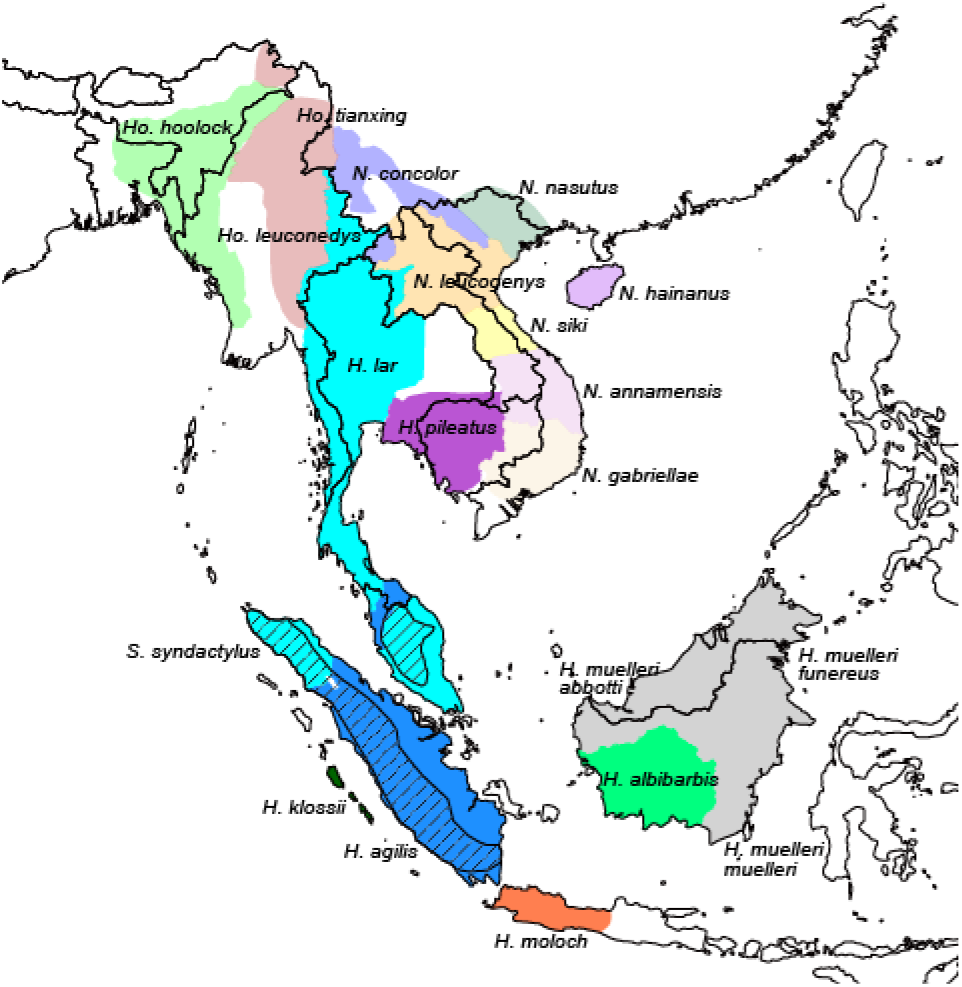
Distribution of extant gibbons. Adapted from Thinh et al. 2010a, 2010b; Fan et al. 2017.

Some studies have proposed to elevate three subspecies of *H. muelleri*, *H. muelleri muelleri* (Müller’s gibbons), *H. muelleri abbotti* (Abbot’s gibbons) and *H. muelleri funereus* (North Bornean grey gibbons), to independent species (Thinh et al. 2010a; Roos 2016); however, no consensus has been reached, thus far. We used the recognised seven-species classification, in this study. In addition, in this study, we refer to *H. pileatus* and *H. lar* as mainland species for simplicity, although some *H. lar* is known to be distributed in northern Sumatra, and refer to the other species as Sundaic species.

The phylogeny of the *Hylobates* species has been studied according to morphology (Groves 1972; Creel and Preuschoft 1984; Geissmann 2002), vocalisation (Geissmann 2002), mitochondrial DNA (mtDNA) (Garza and Woodruff 1992; Hayashi et al. 1995; Takacs et al. 2005; Chatterjee 2006; Whittaker et al. 2007; Chan et al. 2010; Matsudaira and Ishida 2010; Thinh et al. 2010a), and nuclear DNA (Israfil et al. 2011; Kim et al. 2011; Chan et al. 2012, 2013). A whole-mitochondrial genome (mitogenome) study successfully resolved the mtDNA phylogeny of *Hylobates*, which demonstrated that *H. pileatus* diverged first and *H. lar* diverged second, followed by the divergence between proto-*H. agilis*/*H. muelleri* and proto-*H. moloch*/*H. klossii* (Chan et al. 2010 and its correction). This study suggested that divergence began among mainland species and was followed by the Sundaic species, which was supported a scenario in which the common ancestor of *Hylobates* originated on the mainland and that the distribution later expanded to the islands of Southeast Asia (Groves 1972; Chivers 1977; Whittaker et al. 2007). The same and other studies estimated that mtDNA divergence began approximately 3.5 to 3.9 million years ago (MYA) and was completed in a relatively short time (Chan et al. 2010; Matsudaira and Ishida 2010; Thinh et al. 2010a).

In contrast with the mitogenome studies, nuclear DNA markers have yielded ambiguous results on species phylogeny (Kim et al. 2011; Chan et al. 2013), likely due to the rapid divergence that occurred among *Hylobates* species, as suggested by mtDNA, resulting in the incomplete lineage sorting of many nuclear genetic markers. In addition, because hybridisation has been observed in the contact zones between *Hylobates* species (Brockelman and Gittins 1984; Marshall and Sugardjito 1986), many historical hybridisations and introgression may have occurred after the initial divergence of each species; thus, their nuclear genomes may have become admixed. In a previous study, using 14 autosomal loci, covering approximately 11.5 kb, the presence of historical introgression between some *Hylobates* species was detected (Chan et al. 2013). However, the numbers of nuclear DNA markers used in previous studies were likely too few to assess the complicated evolutionary history of the *Hylobates* species. Genome-wide studies that utilise larger numbers of nuclear markers are expected to overcome these potential issues, revealing the detailed evolution of *Hylobates* gibbons. To date, several approaches have been used to conduct genome-wide studies of non-model organisms (Beichman et al. 2018). The whole-genome sequencing (WGS) of several tens of individuals in each species can cover all of the information in the genomes and can account for genetic diversity in each species (e.g., de Manuel et al. 2016; Nater et al. 2017). Although the cost of WGS has been dropping rapidly, studying multiple individuals of many species, such as of *Hylobates*, using WGS remains expensive and is also costly in terms of computational resources.

Reduced-representation sequencing methods, such as restriction-site-associated DNA sequencing (RAD-seq) and genotype by sequencing (GBS), are alternative methods that limit the number of sequencing loci across genomes, facilitating the effective acquisition of sufficient numbers of genetic markers from large numbers of samples, supporting a variety of genome-wide studies (Andrews et al. 2016). Genotyping by random amplicon sequencing-direct (GRAS-Di) is an alternative method that uses polymerase chain reaction (PCR) and a set of primers to amplify several tens of thousands of loci, by randomly annealing the primers to genomes (Enoki and Takeuchi 2018). This method has been successfully applied to a variety of fish species and was able to detect several thousands of single nucleotide variants (SNVs) in each species (Hosoya et al. 2019).

In this study, we conducted a GRAS-Di analysis of gibbon species, to uncover the species phylogeny and the evolutionary history of divergence and hybridisation/introgression events among *Hylobates* gibbons.

## Materials and Methods

### Samples

DNA was extracted from 46 Epstein Barr virus-transformed B lymphoblastoid cell lines and one muscle tissue sample, using the QIAamp DNA Mini Kit (Qiagen, Germany). The DNA samples consisted of two samples from *H. agilis*, three from *H. albibarbis*, 20 from *H. lar*, one from *H. moloch*, four from *H. muelleri* (consisting of two from *H. muelleri muelleri* and two from *H. muelleri abbotti*), 10 from *H. pileatus*, one from *Nomascus leucogenys* (northern white-cheeked gibbons), and six from *Symphalangus syndactylus* (siamangs) (Supplementary Table S1). The samples were derived from captive zoo gibbons, as described in previous studies (Ishida et al. 1985; Nakayama and Ishida 2006).

At the time of sample collection, the species status and key external characteristics were not well-recorded for some individuals. Therefore, we confirmed the maternal species status of all 47 gibbons by comparing the *cytochrome b* (*cytb*) sequence from mtDNA with publicly available data (see details below). To minimise the inclusion of unidentifiable hybrid individuals, we analysed multiple individuals from each species/subspecies, when possible.

### Mitochondrial DNA sequencing and analysis

We determined the *cytb*, tRNA-Thr, tRNA-Pro, and partial D-loop sequences (approximately 1.9 kb) of all 47 gibbons and determined the whole-mitogenome sequences (approximately 16.5 kb) of a subset of individuals (n = 6). Approximately 8-kb (covering the *cytb* to D-loop region) and 10-kb fragments of mtDNA were PCR amplified, using the following primers: 5’-GGCTTTCTCAACTTTTAAAGGATA-3’ and 5’-TGTCCTGATCCAACATCGAG-3’ (8-kb fragment), and 5’-CCGTGCAAAGGTAGCATAATC-3’ and 5’-TTACTTTTATTTGGAGTTGCACCA-3’ (10-kb fragment) (Finstermeier et al. 2013).

PCR was performed in a 25 μl mixture, consisting of 0.5 units KOD FX DNA polymerase (Toyobo, Japan), 1× PCR Buffer for KOD FX, 0.4 mM of each dNTP, 7.5 pmol of each primer, and 1 μl DNA solution. The PCR thermal cycle conditions were as follows: the initial incubation was performed at 94°C, for 2 min, followed by 35 cycles consisted of a denaturation step at 98°C, for 10 s, and an annealing/extension step at 68°C for 8 min (8-kb fragment) or an annealing step at 63°C for 30 s and extension at 68°C for 8 min (10-kb fragment), followed by a final extension step at 68°C for 10 min. PCR amplification was confirmed by agarose gel electrophoresis.

The PCR products were purified using a FavorPrep PCR Clean-Up Mini Kit (Favorgen, Taiwan) and used as templates for sequencing reactions. Sanger sequencing was outsourced to Fasmac (Japan), with internal primers, including three forward primers (5’-CCACGACCAATGATACGAAA-3’; 5’-AGACAACGCCACACTCACAC-3’; and 5’-CTTCACCCTCAGCACCCAAAGC-3’ [Andayani et al. 2001]) and four reverse primers (5’-ATGAGGAGGAGGAGAAATAGTCCTAG-3’; 5’-GGGGTGGAAGGTAATTTTGTC-3’; 5’-CGGCTTACAAGACCGGTGT-3’; and 5’-AAGACAGATACTGCGACATAGG-3’ [Matsudaira et al. 2013]), to determine the *cytb* to D-loop region. Other internal primer sequences are available, on reasonable request, from the authors. Sequences were called from spectrograms, using MEGA 6 (Tamura et al. 2013).

To confirm the maternal species status, we assessed the mtDNA phylogenetic positions of all 47 *cytb* sequences (1,140 bases) by comparing them with the published *cytb* sequences of 175 gibbons (Arnerson et al. 1996; Chan et al. 2010; Matsudaira and Ishida 2010; Thinh et al. 2010a; Thinh et al. 2010b; Finstermeier et al. 2013; Fan et al. 2017) and eight other primate species: *Homo sapiens*, *Pan troglodytes*, *Pan paniscus*, *Gorilla gorilla*, *Pongo pygmaeus*, *Pongo abelli*, *Macaca mulatta*, and *Papio anubis* (Horai et al. 1995; Xu and Arnason 1996; Zinner et al. 2013; Liedigk et al. 2014) (Supplementary Table S2). We constructed a maximum likelihood (ML) tree, using IQ-TREE (Nguyen et al. 2015). The TN+F+I+G4 substitution model was selected by ModelFinder (Kalyaanamoorthy et al. 2017). An ultrafast bootstrap approximation test (Minh et al. 2013) was performed, using 1,000 iterations.

Previous whole-mitogenome studies did not analyse *H. albibarbis* (Chan et al. 2010; Matsudaira and Ishida 2010). Thus, we newly determined the whole-mitogenome sequences of one *H. albibarbis* and five other *Hylobates* gibbons. We analysed these six sequences, together with the published whole-mitogenome sequences of 44 gibbons (which included one *S. syndactylus* in the GRAS-Di data set [Sample ID: G75]) (Arnerson et al. 1996; Chan et al. 2010; Matsudaira and Ishida 2010; Finstermeier et al. 2013; Fan et al. 2017) and eight outgroup species, similar to the *cytb* analysis (Supplementary Table S3). After aligning the sequences using MUSCLE (Edgar 2004), which was implemented in MEGA, we extracted the sequences of 13 protein-coding genes and two rRNA-coding regions, which resulted in 13,798-bp sequences. The ML tree was constructed using IQ-TREE. Among the 15 identified genes, seven partitions with different substitution models were selected by ModelFinder (Supplementary Table S4). An ultrafast bootstrap test was conducted, for 1,000 iterations.

Divergence time estimation was conducted using BEAST 2.6.0. (Bouckaert et al. 2014). The relaxed, lognormal clock model and birth-death model tree priors were selected. The partition scheme and the substitution models used in the ML tree inference were also applied. Three divergence time priors were used, assuming normal distribution: (1) Hominoids and Cercopithecoids, as 30.5 MYA (95% lower and upper limit: 25.0–36.0) (Stevens et al. 2013; Bond et al. 2015), (2) *Pongo* and *Homo*/*Pan*/*Gorilla*, as 15.5 MYA (13.0–18.0) (Kelley 2002; Besenbacher et al. 2019), (3) *Homo* and *Pan*, as 6.5 MYA (6.0–7.0) (Senut et al. 2001; Brunet et al. 2002). Four independent Markov chain Monte Carlo (MCMC) runs were performed for 25,000,000 generations, and parameters were recorded once in every 1,000 generations. The conversions of the results were assessed by Tracer 1.7.1. (Rambaut et al. 2018). The first 25% samples from each run were discarded as burn-in, and the remaining results were merged, using LogCombiner in BEAST. A consensus tree was constructed from 75,004 trees, using TreeAnotator in BEAST, and was visualised using FigTree v1.4.4 (https://github.com/rambaut/figtree/releases).

### GRAS-Di

The RNA in the extracted DNA samples was digested by RNase A (Nippon Gene, Japan), for 1 h. Then, ethanol precipitation was performed, and the DNA was eluted using Low EDTA TE (Thermo Scientific, USA). We measured the DNA concentrations of the samples using a Qubit 4 fluorometer, with the Qubit dsDNA BR Assay Kit (Thermo Scientific), and prepared 100 ng of DNA (> 15 ng/μl) per sample. Library preparation and sequencing using the GRAS-Di method were outsourced to GeneBay, Inc. (Japan). The protocols for the library preparation were those described by Hosoya et al. (2019), except that the PCR reaction volume was 25 μl during the first PCR run and 50 during the second PCR run, instead of 10 μl. The sequences of the 64 primers used to obtain amplicons during the first PCR run were the same as those described in Hosoya et al. (2019). The library was sequenced on a HiSeq 4000 (Illumina, USA), with 151-bp paired-end reads.

### Read mapping and genotyping

Sequence reads were de-multiplexed to each individual and provided as fastq files. We removed 3’ adapter sequences and low-quality reads using cutadapt 1.12 (Martin 2011), with the following command: (cutadapt -a CTGTCTCTTATACACATCTCCGAGCCCACGAGAC -ACTGTCTCTTATACACATCTGACGCTGCCGACGA --overlap 10 --minimum-length 51 --quality-cutoff 20). Cleaned reads were mapped to the reference genome sequence of *N. leucogenys*, nomLeu3 (Carbone et al. 2014), using Burrows-Wheeler Aligner (BWA-MEM) 0.7.17 (Li 2013). Generated sam files were converted into bam files and sorted using SortSam command of Genome Analysis Toolkit (GATK) 4.1.0.0. (McKenna et al. 2010). After adding read group information, using GATK AddOrReplaceReadGroups, each bam file was indexed by samtools 1.9 (Li et al. 2009). GATK HaplotypeCaller with --emit-ref-confidence and GVCF options, was used to call variants in each individual. The generated gvcf files for the 47 gibbons were merged using GATK CombineGVCFs. GATK GenotyepGVCFs, with the -all-sites option, was used for joint genotyping variants among the 47 gibbons.

SNV sites were extracted using GATK SelectVariants. Then, GATK hard filters were applied to remove ambiguous results (QD < 2.0; FS > 60.0; MQ < 40.0; MQRankSum < −12.5; ReadPosRankSum < −8.0). Subsequently, we extracted SNV sites that satisfied the following criteria: (1) genotyped in all 47 gibbons, with at least 5× depth of coverage for each gibbon; (2) biallelic (i.e., no more than two alleles); and (3) variable among the 47 gibbons. Last, we extracted only those SNV sites located on the 25 autosomal long scaffolds of nomLeu3. We used GATK VariantFiltration, vcftools 0.1.17 (Danecek et al. 2011), and vcflib (https://github.com/vcflib/vcflib) to perform the filtering. The generated dataset consisted of 358,142 SNV sites.

### Species tree and network construction

We constructed a phylogenetic tree of the autosomal SNVs, using ML and Bayesian frameworks. The ML tree was constructed using IQ-TREE. Because the dataset consisted of only SNV sites, we used an ascertainment bias correction (ASC) model (Lewis 2001) for the ML tree construction. In the ASC model, heterozygous and invariant sites were ignored; thus, 196,186 of 358,142 SNV sites were used for the analysis. The TVM+F+ASC+R2 model was selected by ModelFinder, and an ultrafast bootstrap approximation test was performed, with 1,000 iterations.

A Bayesian tree was constructed, using MrBayes 3.2.7a (Ronquist et al. 2012). The GTR+G model was selected by MrModeltest 2.4 (Nylander 2004). Four independent MCMC runs were performed, consisting of one heat chain and three cold chains. Each run comprised 10,000,000 generations, and the parameters were sampled every 5,000 generations. Convergence among the four runs was assessed by Tracer. The first 25% of samples from each run were discarded as burn-in, and the results of the four runs (i.e., 6,004 samples) were merged. The tree was visualised using FigTree.

To assess the reticulate evolution among *Hylobates*, a phylogenetic network of autosomal SNVs was generated, using the NeighborNet algorithm in SplitsTree 4.15.1 (Huson and Bryant 2006). To clarify the results, we only included *Hylobates* gibbons in this analysis.

### Allele-sharing distance

To uncover the genetic relationships among individuals, we calculated the allele-sharing distances (ASD) (Bowcock et al. 1994) between each pair of individuals, using asd v1.0.0 (https://github.com/szpiech/asd). Based on the ASD matrix, multi-dimensional scaling (MDS) plots (two-dimensional plots) were drawn, using R 3.6.2 (R Core Team 2019).

### Detection of introgression

The presence of introgression among *Hylobates* was assessed by Patterson’s D-statistics tests (also known as ABBA-BABA tests) (Green et al. 2010; Patterson et al. 2012). Because the phylogenetic relationships among *Hylobates* species/subspecies have been inferred from both mitogenomes and autosomal SNVs and are well-supported and consistent, (except for one mitogenome from *H. agilis*; see Results), we used the autosomal tree topology and calculated the D-statistics by setting the outgroup as either *N. leucogenys* or *S. syndactylus*. The D-statistics were calculated for both combinations of one individual per species/subspecies and the combinations of population data for species/subspecies. The D-statistics were calculated using AdmixTools (Patterson et al. 2012), with the R package admixr (Petr et al. 2019).

### Potentially introgressed loci

When an introgression event occurs, most genomic regions from other species are expected to be removed, due to incompatibility and deleteriousness. However, some regions may remain as introgressed loci because of adaptive advantages. Potentially introgressed regions can be detected by comparing D-statistics and genetic divergence in each segment of the genome (Smith and Kronfrost 2013). Because introgressed regions should have shorter coalescent times than average, the regions with high absolute D-statistics and low genetic divergence can be considered candidates for introgressed regions.

To detect the potentially introgressed loci among *Hylobates* species/subspecies, we calculated the corrected D-statistics, *f_d_*, (Martin et al. 2015), and genetic divergence *d_xy_* (Smith and Kronfrost 2013), using ABBABABAwindows.py and popgenWindows.py (https://github.com/simonhmartin/genomics_general), respectively. Because of limitations in the number of SNV sites in our data set, we calculated the values for regions with a large window size of 1,000,000 bases, with at least 100 SNVs per window.

## Results

### Genotyping

We obtained a mean ± standard deviation (SD) of 8.6 million (M) ± 0.87 (min = 6.6 M; max = 10.2 M) raw read pairs, per sample. Adapter trimming and quality filtering by cutadapt yielded 8.5 M ± 0.85 (min = 6.5 M; max = 10.0 M) read pairs, per sample. In the nomLeu3 genome, 3.6% to 5.2% (95,030,378 to 139,035,756 sites) of the 2,654,007,897 autosomal sites were mapped by at least one read in each sample, and 1.2% to 1.8% (31,448,837 to 47,600,791 sites) were mapped by at least five reads in each sample. The mean depth of coverage at the mapping sites in each sample was 16.3× ± 1.4. After hard filtering by GATK, a data set containing 358,142 biallelic, autosomal, SNV sites that were successfully genotyped in all 47 gibbons was generated.

We calculated the nucleotide diversity (Nei and Li 1979) of each species/subspecies across the 358,142 autosomal SNV sites (Table 1). *H. pileatus* showed the lowest nucleotide diversity (0.0198 ± 0.0811). The three Bornean species/subspecies (*H. albibarbis*, *H. muelleri muelleri*, and *H. muelleri abbotti*) showed the highest nucleotide diversity (0.0598–0.0694). The other three *Hylobates* species (*H. moloch*, *H. lar*, and *H. agilis*), *S. syndactylus*, and *N. leucogenys* were in the middle (0.0371–0.0511).

**Table 1.** Nucleotide diversity (π) across the 358 142 SNV sites.

| Species | Sample size | $\pi$ (± SD) |
| --- | --- | --- |
| *H. pileatus* | 10 | 0.0198 ± 0.0811 |
| *H. lar* | 20 | 0.0489 ± 0.1240 |
| *H. moloch* | 1 | 0.0371 ± 0.1891 |
| *H. agilis* | 2 | 0.0511 ± 0.1605 |
| *H. albibarbis* | 3 | 0.0646 ± 0.1543 |
| *H. muelleri muelleri* | 2 | 0.0598 ± 0.1718 |
| *H. muelleri abboti* | 2 | 0.0694 ± 0.1814 |
| *S. syndactylus* | 6 | 0.0415 ± 0.1151 |
| *N. leucogenys* | 1 | 0.0462 ± 0.2100 |

### Phylogenetic tree, divergence time, ASD, and phylogenetic network

Phylogenetic relationships among *Hylobates* were well resolved, both in the mitogenome and the autosomal SNV trees (Fig. 2), and the topologies of species/subspecies relationships were consistent, except for one mitogenome sequence from *H. agilis* (Sample ID: G80). This individual clustered with the other *H. agilis* individuals in the autosomal SNV tree (Fig. 2B) but clustered with *H. lar* individuals and separated from other *H. agilis* individuals in the mitogenome tree (Fig. 2A; labelled as *H. agilis* [group2]). In the *cytb* tree, which consisted of the reference sequences of four *H. lar* subspecies, this *H. agilis* individual tightly clustered with the sequences from *H. lar vestitus* (Sumatran white-handed gibbons) (Supplementary Fig. S1). Because the mitogenome represents the genealogy of only a single locus, we considered the autosomal SNV tree to represent the true species tree and used the autosomal SNV tree for all further analyses.

**Fig. 2.**
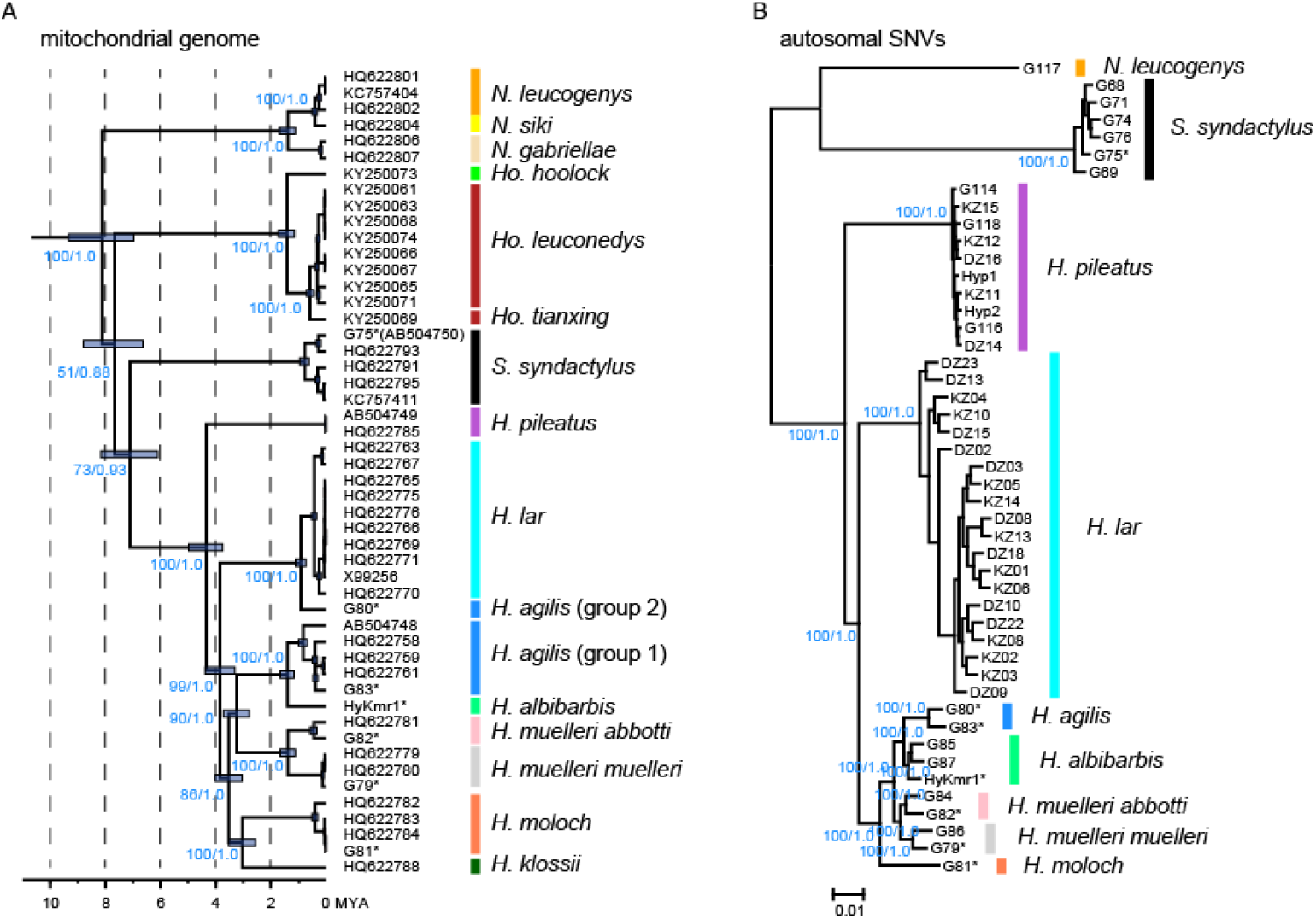
Phylogenetic trees of gibbons. Values on the nodes (shown in blue) are bootstrap values for the ML analysis and the posterior probabilities for the Bayesian analysis. (A) Mitogenome phylogeny and divergence times, as estimated by Bayesian analysis, using BEAST 2. The outgroup species were omitted from this figure. (B) Species phylogeny, based on the autosomal SNVs, produced by ML analysis using IQ-TREE. * indicates the individuals (n = 7) that were analysed both for mitogenomes and autosomal SNVs.

Both the autosomal and mitogenome phylogenetic trees demonstrated that the divergence of *Hylobates* species started in the mainland species, followed by the Sundaic species/subspecies. Based on the estimated divergence time indicated by the mitogenomes (Fig. 2A, Supplementary Table S5), among *Hylobates*, divergence first occurred between *H. pileatus* and the common ancestor of the other *Hylobates* species, approximately 4.3 MYA (95% highest posterior density credibility interval [HPD CI]: 3.7 to 5.0). Next, *H. lar* diverged from the Sundaic species, approximately 3.8 MYA (3.3 to 4.4). Then, *H. moloch* (and *H. klossii*) diverged from the Sumatran/Bornean species, approximately 3.5 MYA (3.0 to 4.0). *H. muelleri* and *H. agilis*/*H. albibarbis* diverged approximately 3.2 MYA (2.8 to 3.7). *H. agilis* and *H. albibarbis* diverged approximately 1.4 MYA (1.2 to 1.7). *H. muelleri muelleri* and *H. muelleri abbotti* diverged approximately 1.4 MYA (1.1 to 1.7).

MDS plots of ASD also revealed similar relationships among the *Hylobates* species/subspecies. In the MDS plot containing all *Hylobates* gibbons (n = 40), three clusters appeared. *H. pileatus* formed one cluster, *H. lar* formed another cluster, and the Sundaic *Hylobates* species/subspecies formed the final cluster (Fig. 3A). In addition, among the Sundaic *Hylobates* species/subspecies, *H. moloch* separated from the others, which is consistent with the autosomal and mitogenome phylogeny results. When we plotted only the Sundaic *Hylobates* species/subspecies (n = 10), the species/subspecies were all separated from each other (Fig. 3B). *H. agilis*, *H. albibarbis*, and *H. muelleri* (*muelleri* and *abbotti* subspecies) were plotted on a line, with *H. agilis* at one end and *H. muelleri* at the other end. *H. albibarbis* was located between the two species, closer to *H. agilis* than to *H. muelleri*. Furthermore, *H. muelleri muelleri* (n = 2) and *H. muelleri abbotti* (n = 2) were slightly separated from each other in the plot, which indicated the presence of genetic structure-divergence between the two subspecies.

**Fig. 3.**
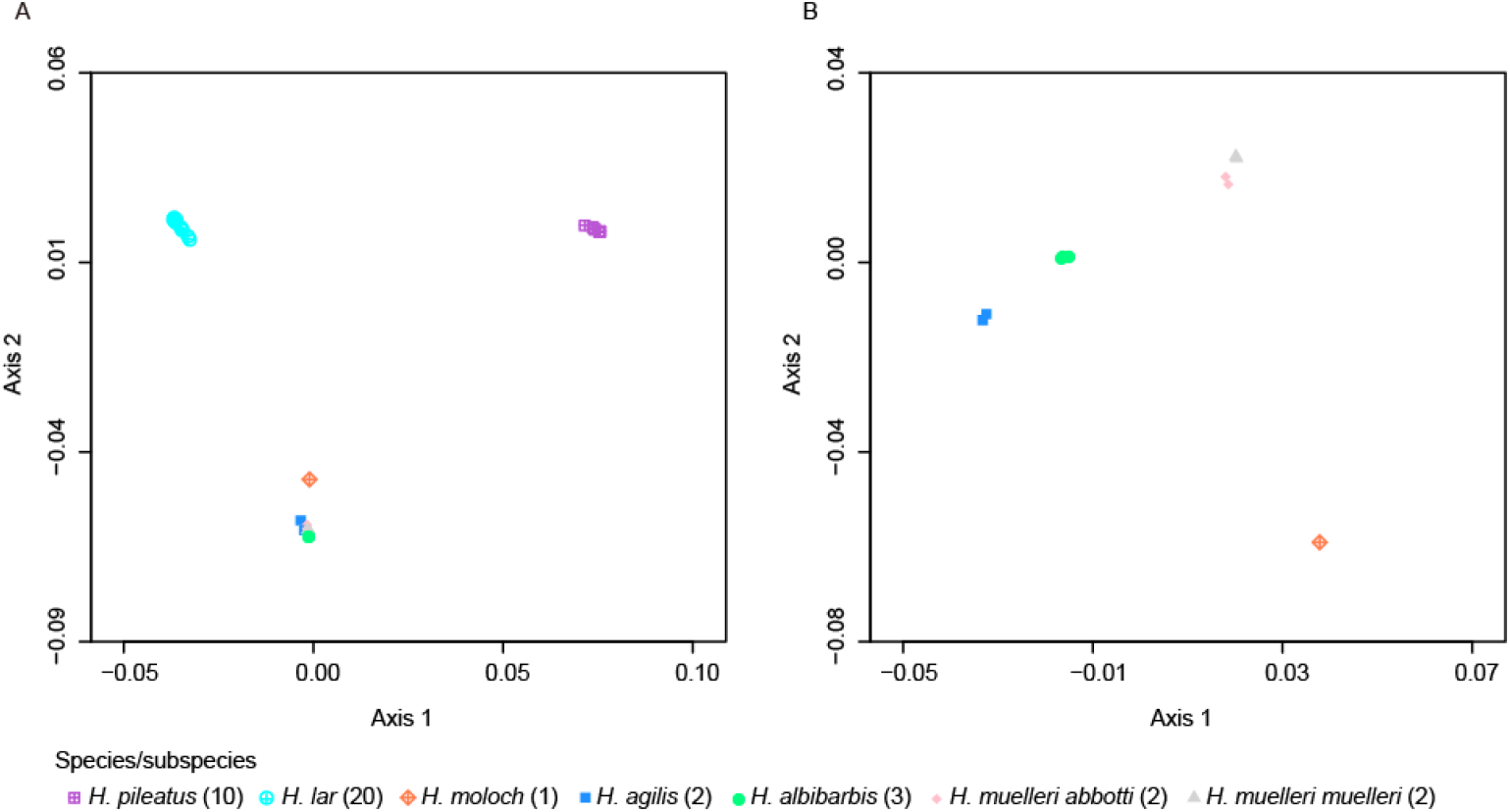
MDS plot of ASD. (A) A two-dimensional plot of ASD for seven *Hylobates* species/subspecies (n = 40). (B) A two-dimensional plot of ASD for five Sundaic *Hylobates* species/subspecies (n = 10).

The phylogenetic network (Supplementary Fig. S2) showed consistent relationships among the *Hylobates* species/subspecies, as shown in the phylogenetic trees and the MDS plots of ASD. A clear reticulation, which indicates the introgression, was observed in *H. albibarbis* and *H. muelleri* (both *muelleri* and *abbotti* subspecies). In addition, the two *H. agilis* individuals (Sample ID: G80 and G83) formed a reticulation, which may be related to differences in their mitogenomes (Fig. 2A). Furthermore, the network highlighted the presence of a population structure within *H. lar*, in which 14 of 20 individuals were tightly clustered together. This result was similar to the *cytb* tree, in which the same 14 individuals and one additional individual were clustered more tightly compared with the other five individuals (Supplementary Fig. S1).

### Introgression

The D-statistics calculated for species quartets showed signals of introgression (i.e., absolute Z-score > 3) between *H. albibarbis* and *H. muelleri* (both *muelleri* and *abbotti* subspecies), between *H. pileatus* and *H. lar*, between *H. lar* and *H. agilis*, between *H. albibarbis* and *H. moloch*, and between *H. muelleri abbotti* and *H. moloch* (Table 2, Supplementary Table S6). These results were consistent, regardless of the outgroup species (*N. leucogenys* and *S. syndactylus*) used for the analysis. No other signals of introgression were detected among the other testable quartets of *Hylobates* species/subspecies. The D-statistics calculated for individual quartets also showed signals of introgression in the same pairs of species/subspecies (and several cases between the *H. muelleri abbotti* and *H. lar* individuals) (Fig. 4, Supplementary Fig. S3, Supplementary Table S7). However, the differences in the genetic makeups of the two *H. agilis* individuals (Sample ID: G80 and G83) affected the D-statistics and Z-scores, and the signals could be both higher and lower than the threshold (absolute Z-score > 3), in some cases, such as between *H. albibarbis* and *H. moloch*.

**Table 2.**
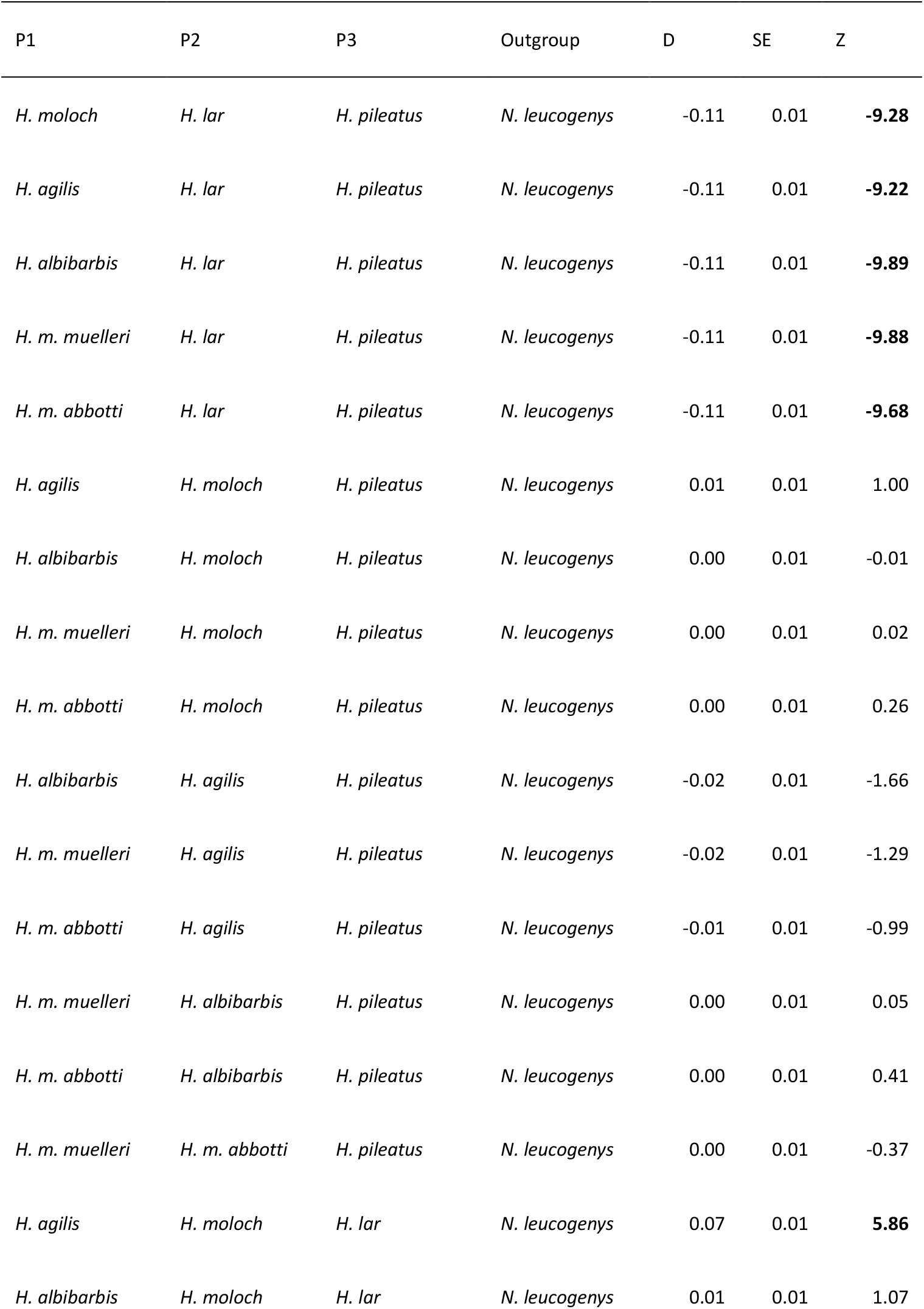
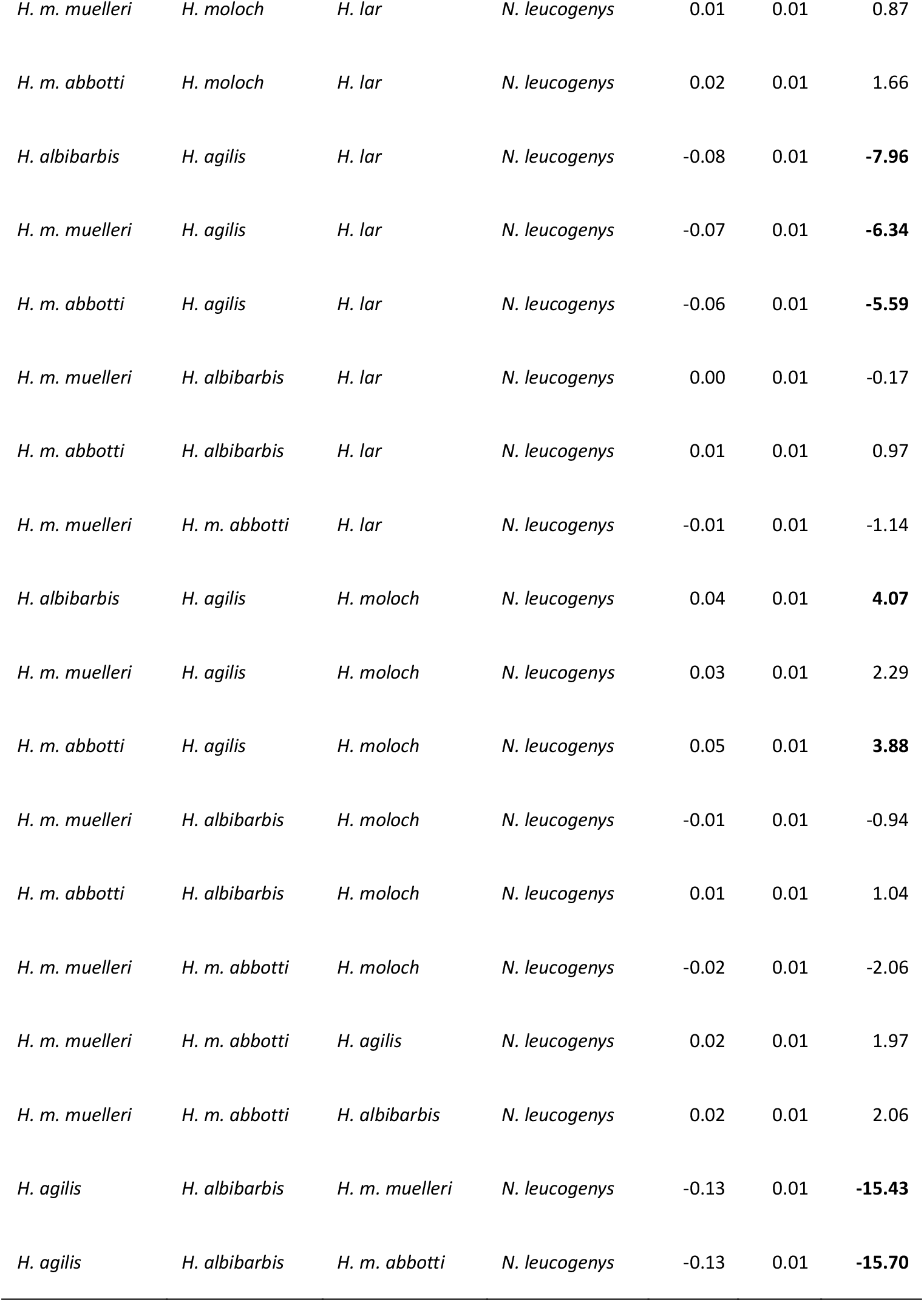
Patterson’s D-statistics by species. *N. leucogenys* was used as an outgroup. Positive Z-scores (> 3) indicate the introgression between P1 and P3. Negative Z-scores (< −3) indicate the introgression between P2 and P3.

| P1 | P2 | P3 | Outgroup | D | SE | Z |
| --- | --- | --- | --- | --- | --- | --- |
| *H. moloch* | *H. lar* | *H. pileatus* | *N. leucogenys* | -0.11 | 0.01 | **-9.28** |
| *H. agilis* | *H. lar* | *H. pileatus* | *N. leucogenys* | -0.11 | 0.01 | **-9.22** |
| *H. albibarbis* | *H. lar* | *H. pileatus* | *N. leucogenys* | -0.11 | 0.01 | **-9.89** |
| *H. m. muelleri* | *H. lar* | *H. pileatus* | *N. leucogenys* | -0.11 | 0.01 | **-9.88** |
| *H. m. abbotti* | *H. lar* | *H. pileatus* | *N. leucogenys* | -0.11 | 0.01 | **-9.68** |
| *H. agilis* | *H. moloch* | *H. pileatus* | *N. leucogenys* | 0.01 | 0.01 | 1.00 |
| *H. albibarbis* | *H. moloch* | *H. pileatus* | *N. leucogenys* | 0.00 | 0.01 | -0.01 |
| *H. m. muelleri* | *H. moloch* | *H. pileatus* | *N. leucogenys* | 0.00 | 0.01 | 0.02 |
| *H. m. abbotti* | *H. moloch* | *H. pileatus* | *N. leucogenys* | 0.00 | 0.01 | 0.26 |
| *H. albibarbis* | *H. agilis* | *H. pileatus* | *N. leucogenys* | -0.02 | 0.01 | -1.66 |
| *H. m. muelleri* | *H. agilis* | *H. pileatus* | *N. leucogenys* | -0.02 | 0.01 | -1.29 |
| *H. m. abbotti* | *H. agilis* | *H. pileatus* | *N. leucogenys* | -0.01 | 0.01 | -0.99 |
| *H. m. muelleri* | *H. albibarbis* | *H. pileatus* | *N. leucogenys* | 0.00 | 0.01 | 0.05 |
| *H. m. abbotti* | *H. albibarbis* | *H. pileatus* | *N. leucogenys* | 0.00 | 0.01 | 0.41 |
| *H. m. muelleri* | *H. m. abbotti* | *H. pileatus* | *N. leucogenys* | 0.00 | 0.01 | -0.37 |
| *H. agilis* | *H. moloch* | *H. lar* | *N. leucogenys* | 0.07 | 0.01 | **5.86** |
| *H. albibarbis* | *H. moloch* | *H. lar* | *N. leucogenys* | 0.01 | 0.01 | 1.07 |
| *H. m. muelleri* | *H. moloch* | *H. lar* | *N. leucogenys* | 0.01 | 0.01 | 0.87 |
| *H. m. abbotti* | *H. moloch* | *H. lar* | *N. leucogenys* | 0.02 | 0.01 | 1.66 |
| *H. albibarbis* | *H. agilis* | *H. lar* | *N. leucogenys* | -0.08 | 0.01 | **-7.96** |
| *H. m. muelleri* | *H. agilis* | *H. lar* | *N. leucogenys* | -0.07 | 0.01 | **-6.34** |
| *H. m. abbotti* | *H. agilis* | *H. lar* | *N. leucogenys* | -0.06 | 0.01 | **-5.59** |
| *H. m. muelleri* | *H. albibarbis* | *H. lar* | *N. leucogenys* | 0.00 | 0.01 | -0.17 |
| *H. m. abbotti* | *H. albibarbis* | *H. lar* | *N. leucogenys* | 0.01 | 0.01 | 0.97 |
| *H. m. muelleri* | *H. m. abbotti* | *H. lar* | *N. leucogenys* | -0.01 | 0.01 | -1.14 |
| *H. albibarbis* | *H. agilis* | *H. moloch* | *N. leucogenys* | 0.04 | 0.01 | **4.07** |
| *H. m. muelleri* | *H. agilis* | *H. moloch* | *N. leucogenys* | 0.03 | 0.01 | 2.29 |
| *H. m. abbotti* | *H. agilis* | *H. moloch* | *N. leucogenys* | 0.05 | 0.01 | **3.88** |
| *H. m. muelleri* | *H. albibarbis* | *H. moloch* | *N. leucogenys* | -0.01 | 0.01 | -0.94 |
| *H. m. abbotti* | *H. albibarbis* | *H. moloch* | *N. leucogenys* | 0.01 | 0.01 | 1.04 |
| *H. m. muelleri* | *H. m. abbotti* | *H. moloch* | *N. leucogenys* | -0.02 | 0.01 | -2.06 |
| *H. m. muelleri* | *H. m. abbotti* | *H. agilis* | *N. leucogenys* | 0.02 | 0.01 | 1.97 |
| *H. m. muelleri* | *H. m. abbotti* | *H. albibarbis* | *N. leucogenys* | 0.02 | 0.01 | 2.06 |
| *H. agilis* | *H. albibarbis* | *H. m. muelleri* | *N. leucogenys* | -0.13 | 0.01 | **-15.43** |
| *H. agilis* | *H. albibarbis* | *H. m. abbotti* | *N. leucogenys* | -0.13 | 0.01 | **-15.70** |

**Fig. 4.**
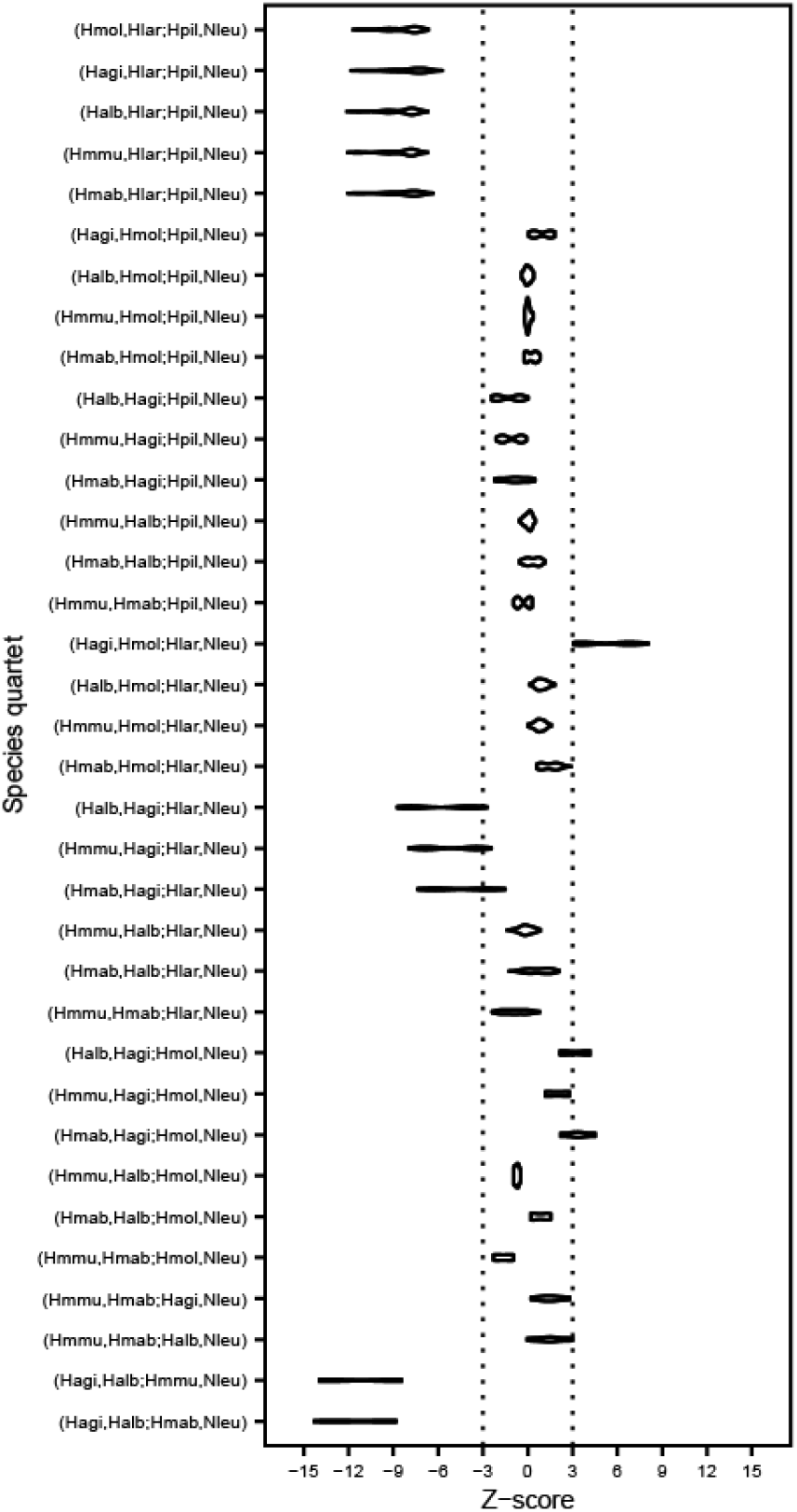
The ranges of Z-scores for Patterson’s D-statistics among *Hylobates* species/subspecies. *N. leucogenys* was using as an outgroup. Species quartets are shown as (P1, P2; P3, Outgroup). Positive Z-scores (> 3) indicate introgression between P1 and P3. Negative Z-scores (< −3) indicate introgression between P2 and P3. Hagi: *H. agilis*, Halb: *H. albibarbis*, Hlar: *H. lar*, Hmab: *H. muelleri abbotti*, Hmmu: *H. muelleri muelleri*, Hmol: *H. moloch*, Hpil: *H. pileatus*, Nleu: *N. leucogenys*.

### Potentially introgressed loci

Calculations of *f_d_* and *d_xy_* were performed for the pairs of species whose introgression was detected by D-statistics. We considered autosomal regions with the top 5% *f_d_* values and the bottom 5% *d_xy_* values to be potentially introgressed loci (Supplementary Table S8). Among those, three loci (chr7b: 50,000,001-51,000,000; chr7b: 51,000,0001–52,000,000; and chr10: 68,000,0001-69,000,000) were detected as candidates between *H. lar* and *H. pileatus*, in all of the tested quartets (Fig. 5, Supplementary Table S8). Furthermore, two loci (chr4: 151,000,001–152,000,000; chr16: 55,000,001–56,000,000) were detected between *H. agilis* and *H. lar*, in several tested quartets (Supplementary Table S8). Note that the genomic positions shown here are those for *N. leucogenys* (2n = 52) and, thus, differ from the exact genomic positions found in *Hylobates* (2n = 44).

**Fig. 5.**
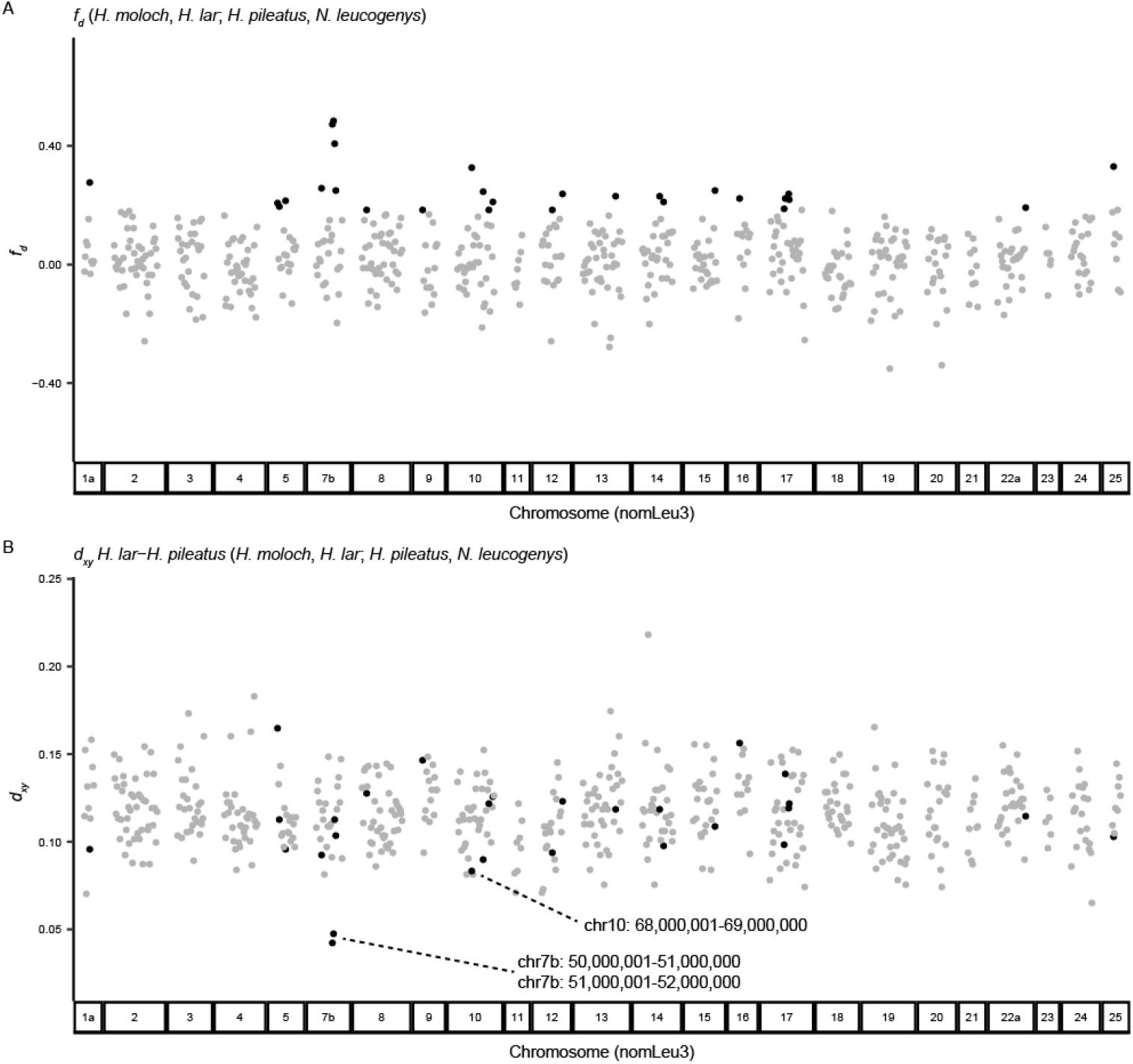
Potentially introgressed genomic regions between *H. lar* and *H. pileatus*. (A) *f_d_* (Martin et al. 2015) and (B) *d_xy_* were calculated for 1 M base-windows, containing more than 100 SNV sites. Genomic regions with the top 5% of *f_d_* values are shown as black dots, whereas all others are shown as grey dots. The genomic regions are presented using the reference genome of *N. leucogenys* (nomLeu3; 2n = 52) and, thus, do not match the chromosomal conditions of *Hylobates* (2n = 44).

## Discussion

The autosomal phylogeny identified in this study was consistent with the previously reported mitogenome phylogeny (Chan et al. 2010 and its correction) and the mitogenome phylogeny identified in this study, which supports the following scenario: the most recent common ancestor of *Hylobates* originated on the mainland in Southeast Asia, expanded their distribution southward and emerged in Sundaland during the Pliocene era (approximately 3.0 to 4.3 MYA), and speciated due to subsequent isolations (Whittaker et al. 2007; Thinh et al. 2010a; Chan et al. 2013).

The early divergence of *H. moloch* on Java, relative to the other Sundaic species living in Sumatra and Borneo, which was previously uncovered by the earlier mitogenome study (Chan et al. 2010) was also supported by the present autosomal phylogeny study. We could not include *H. klossii* on the Mentawai islands in the GRAS-Di analysis; thus, the close relationship between *H. moloch* and *H. klossii* that has been suggested by mtDNA phylogeny (Takacs et al. 2005; Chan et al. 2010) and vocalisation analysis (Geissmann 2002) should be tested in the future.

Both the mitogenome and the autosomal trees suggested that soon after the divergence of *H. moloch*, *H. agilis*/*H. albibarbis* and *H. muelleri* diverged from each other. ASD also suggests that *H. albibarbis* has a closer relationship with *H. agilis* than with *H. muelleri* (*muelleri* and *abbotti* subspecies). This result is consistent with a previous population-based genetic study (Hirai et al. 2009), a *cytb* phylogeny analysis (Thinh et al. 2010a), and a previous taxonomy classification, which included *H. albibarbis* as a subspecies of *H. agilis*, based on vocalisations (Marshall and Sugardjito 1986).

At present, *H. albibarbis* and *H. muelleri* are distributed in Borneo, whereas *H. agilis* is distributed in Sumatra. The current distribution may have occurred as follows; the common ancestor of *H. agilis* and *H. albibarbis* was first distributed only in Sumatra, and approximately 1.4 MYA, the proto-*H. albibarbis* extended their distribution to southwestern Borneo and either replaced or admixed with *Hylobates* gibbons (likely *H. muelleri*) that once lived in that region (Thinh et al. 2010a). Lower sea levels and the appearance of land bridges in the Sundaic region during glacial periods (Woodruff 2010) likely made the migration between Sumatra and Borneo possible.

Alternatively, the common ancestor of *H. agilis* and *H. albibarbis* may have widely distributed, from Sumatra to southwestern Borneo, after the divergence from proto-*H. muelleri*, approximately 3.2 MYA, and 1.4 MYA may represent the time point at which the forests of Sumatra and southwestern Borneo became disconnected. Interestingly, the estimated mtDNA divergence time between *H. muelleri muelleri* and *H. muelleri abbotti* was also approximately 1.4 MYA. Thus, the fragmentation of forests and the isolation of gibbon populations in the Sundaic region may have become more significant during this period than before.

Both the mtDNA divergence time and the ASD of autosomal SNVs between *H. muelleri muelleri* and *H. muelleri abbotti* (0.0697 to 0.0701; n = 4 pairs) were similar to (or slightly larger than) those between *H. agilis* and *H. albibarbis* (0.0671–0.0693; n = 6 pairs). Therefore, these results suggest that these two taxa may be better classified as different species (i.e., *H. muelleri* and *H. abbotti*), which was proposed by Thinh et al. (2010a). However, these results are not conclusive, because the sample size of this study was limited and only two individuals in each subspecies were analysed. In addition, *H. muelleri funereus* was not analysed in this study. The analysis of a larger number of samples, including *H. muelleri funereus*, remains necessary to determine whether our results successfully captured the genetic diversity among *H. muelleri* and to draw conclusions regarding their taxonomic statuses.

The GRAS-Di analysis detected introgression signals between several pairs of *Hylobates* species/subspecies in Patterson’s D-statistics. The signals of introgression between *H. lar* and *H. pileatus*, between *H. albibarbis* and *H. muelleri* (both *muelleri* and *abbotti* subspecies), and between *H. lar* and *H. agilis* were detected in every individual of each species. Interestingly, these three species pairs have been known to form contact/hybrid zones at the species boundaries (Brockelman and Gittins 1984); however, at present, the contact/hybrid zones are limited to small areas. Because every individual of the species demonstrated signals of introgression, the limited hybridisation and introgression that has recently been observed in the contact/hybrid zones appears insufficient to explain the results. Following the glacial cycles that formed the Sundaland and fragmented the forests (Woodruff 2010), the shapes of species boundaries and hybrid zones are likely to have changed over time.

The degree of admixed ancestry that can be identified in individuals from the same species/subspecies may not be uniform across their distribution. Two individuals of *H. agilis* (Sample ID: G80 and G83) showed different levels of D-statistic and Z-score values in several combinations, which likely reflects differences in the amounts of *H. lar*-derived genetic components in their genetic makeups. Individual G80, which has a mitogenome that is closely related to *H. lar* (*vestitus* subspecies), showed higher absolute D-statistics in tests with *H. lar*. This type of mtDNA has been observed in several *H. agilis* individuals from Sumatra, in a previous study (Hirai et al. 2009). Thus, the mtDNA of G80 likely introgressed from *H. lar* to *H. agilis* in the past, similar to the observed mtDNA introgression from *H. pileatus* to *H. lar* (Matsudaira et al. 2013). The frequency and distribution of this mtDNA group among *H. agilis* are not known, but the differences in the mtDNA and the D-statistics may reflect the distance from the species boundary between *H. lar* and *H. agilis*, which is located around Lake Toba. Because differences in the genetic makeup between G80 and G83 affected the D-statistic values, future studies of *Hylobates* gibbons should be aware of potential hidden genetic structures within a species.

Although the introgression signals were not robust among every individual, introgression between *H. albibarbis* and *H. moloch* and between *H. muelleri abbotti* and *H. moloch* were also suggested. For these two species pairs, the current distributions are separated by seas; however, these distributions may have contacted each other on land bridges, when the sea level was low. These results suggested that whenever the distribution range of two *Hylobates* species formed contact regions, hybrid zones were formed and that the reproductive barriers between *Hylobates* species, other than geographical barriers, are incomplete. However, the degrees of introgression may vary among hybrid zones, as suggested by previous observations. Although the degree of introgression may be limited between *H. lar* and *H. pileatus* within the hybrid zone at Khao Yai, Thailand (Brockelman and Gittins 1984), a hybrid swarm (a stable hybrid/admixed population) has formed between *H. albibarbis* and *H. muelleri* within a hybrid zone at the Barito headwaters, central Borneo (Mather 1992).

Previous studies have suggested possible factors that may limit hybridisation and introgression in these contact/hybrid zones, such as positive assortative mating and the reduced fitness of hybrid gibbons (Brockelman and Gittins 1984, Mather 1992, Suwanvecho and Brockelman 2012; Asensio et al. 2017). Further genetic studies that focus on such hybrid zones are expected to reveal factors that affect the frequency of hybridisation and the degree of introgression among species.

Some potentially introgressed regions were detected by assessing their *f_d_* and *d_xy_* values; however, this analysis was unable to detect particular genes or sets of introgressed SNVs because the density of SNVs available in this study was low; thus, we were only able to conduct the test using large window sizes. Intensive sequencing efforts, such as the WGS of *Hylobates* species, will reveal additional details regarding the introgressed regions detected in this study, and will likely detect additional regions across the genome.

Overall, the GRAS-Di analysis, which successfully identified hundreds of thousands of autosomal SNVs, effectively resolved the species phylogeny of *Hylobates* gibbons (except *H. klossii*) and revealed the presence of hybridisation and introgression throughout the evolution of *Hylobates*. Importantly, this study suggested that the phenotypic diversity of *Hylobates* gibbons that is currently observed, such as in pelage colour and vocalisation, represent the products of both divergence and hybridisation/introgression. This study supports the current view of hybridisation/introgression as driving factors in the evolution of primates and other taxa. Future studies of *Hylobates* gibbons that include more samples with known origins and WGS studies will likely uncover additional details regarding the contributions of genetic changes that have occurred during divergence and introgression to species/subspecies differences and diversity among *Hylobates*.

## Acknowledgements

This study was supported by JSPS KAKENHI [Grant Numbers JP18H02516 and JP19K16240].

## Data archiving

The raw read files produced by the GRAS-Di analysis were deposited in DDBJ Sequence Read Archive (DRA), with BioProject Accession No. PRJDB9913. The mtDNA sequences were deposited in the DDBJ/EMBL/NCBI nucleotide database, with Accession No. LC548011-LC548056. Please see Supplementary Table S1 for the details.

