## Supplementary Figures 1-3 for "Divergence and introgression in small apes, the genus *Hylobates*, revealed by reduced representation sequencing"

Fig. S1 Mitochondrial DNA-based ML phylogenetic tree, based on the *cytb* sequences (1,140 bases).
Fig. S2 NeighborNet phylogenetic network of *Hylobates* gibbons based on the autosomal SNVs.
Fig. S3 The ranges of Z-scores for Patterson’s D-statistics among *Hylobates* species/subspecies.

Table S1 List of samples and deposited data information.
Table S2 List of the *cytb* sequences from the database.
Table S3 List of mitochondrial genome sequences from the database.
Table S4 Partition scheme and substitution models used for the mitochondrial genome analysis.
Table S5 Estimated divergence time of mitochondrial genomes.
Table S6 Patterson's D-statistics by species.
Table S7 Patterson's D-statistics by individuals.
Table S8 Potentially introgressed loci.

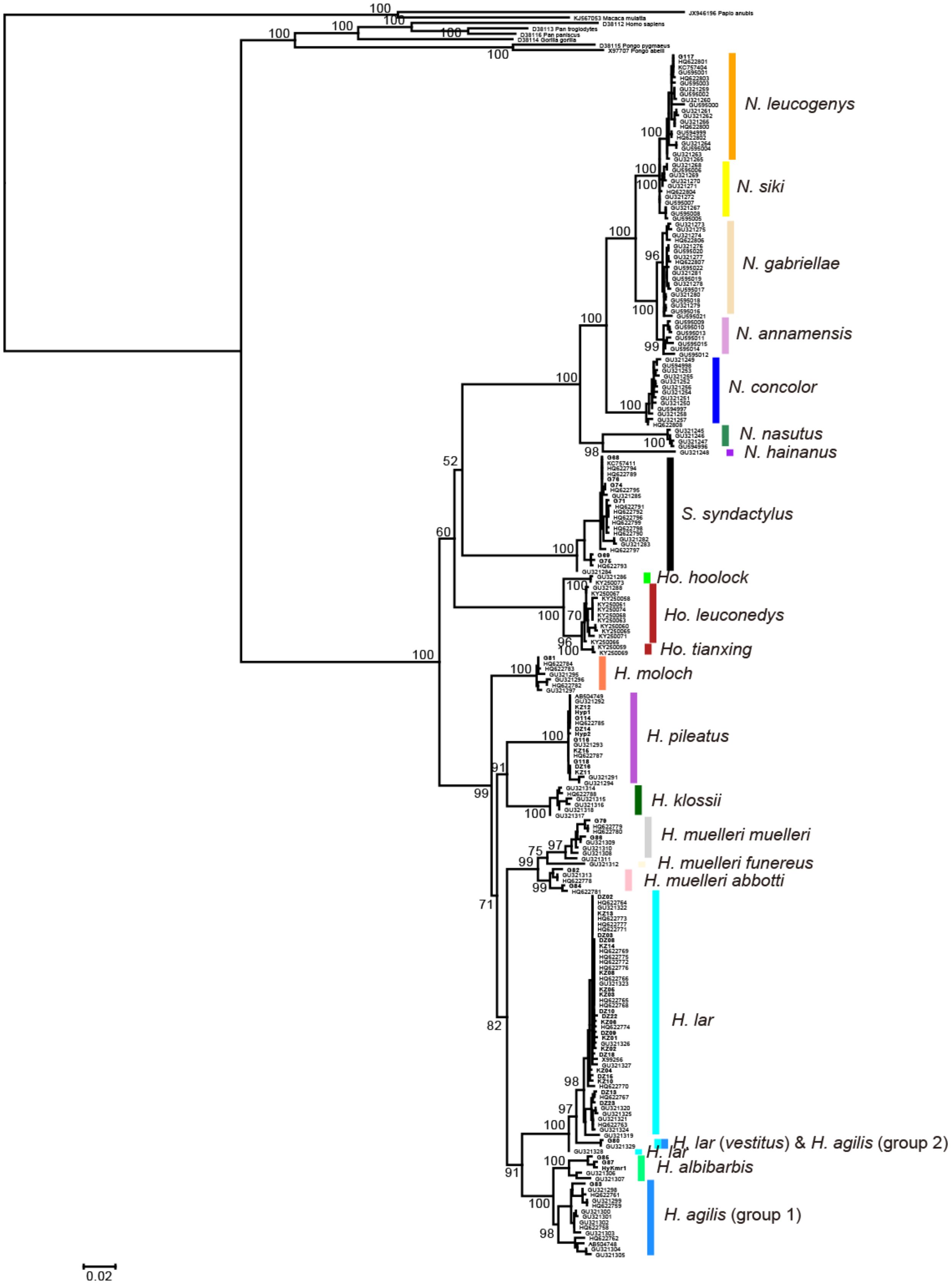


**Fig. S1** Mitochondrial DNA-based ML phylogenetic tree, based on the *cytb* sequences (1,140 bases). Values on nodes are bootstrap values.

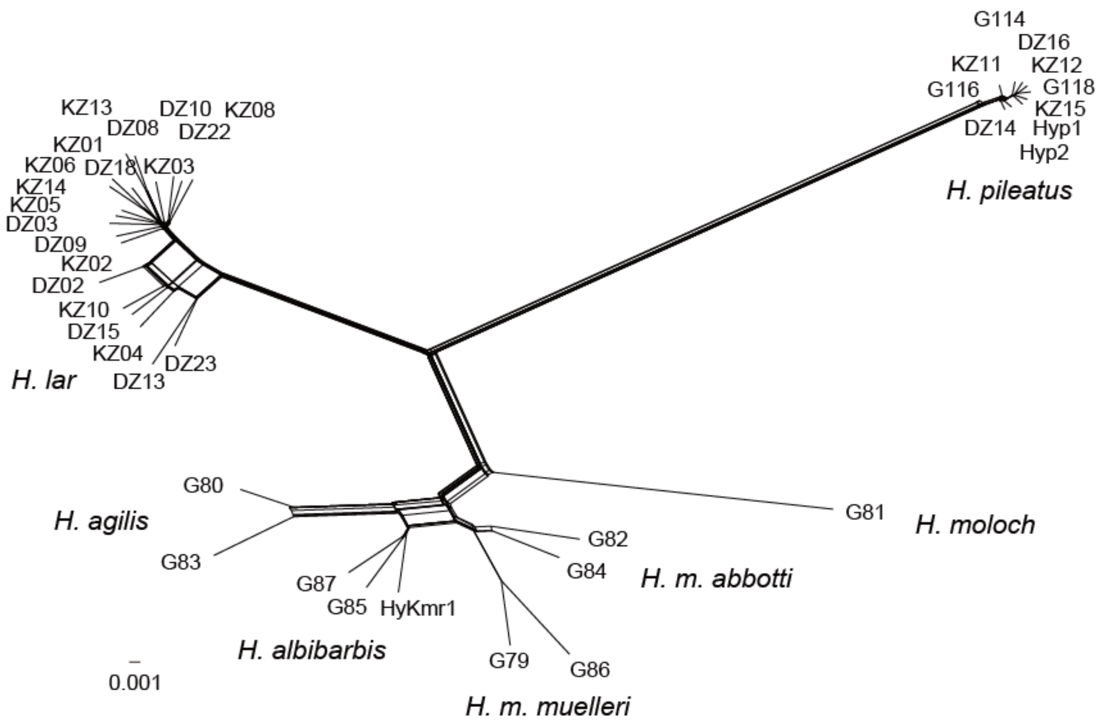


**Fig. S2** NeighborNet phylogenetic network of *Hylobates* gibbons based on the autosomal SNVs.

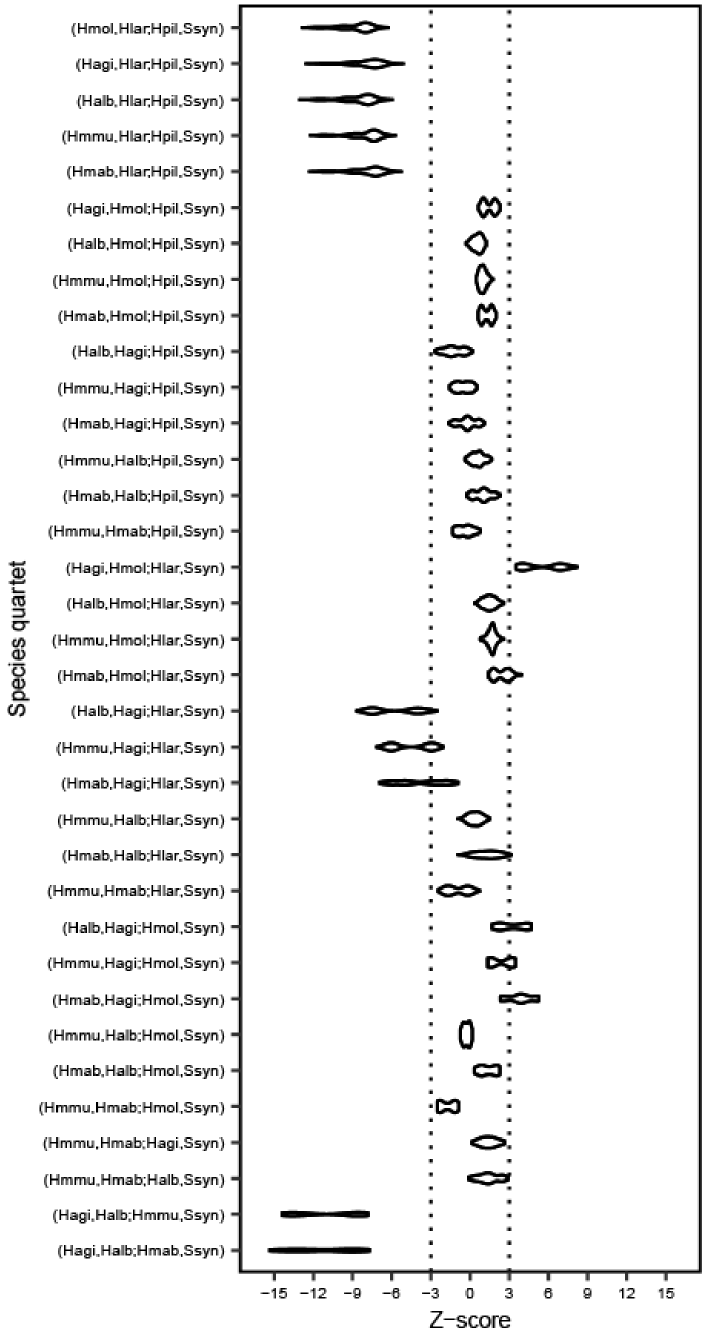


**Fig. S3** The ranges of Z-scores for Patterson's D-statistics among *Hylobates* species/subspecies. *S. syndactylus* was using as an outgroup. Species quartets are shown as (P1, P2; P3, Outgroup). Positive Z-scores (> 3) indicate introgression between P1 and P3. Negative Z-scores (< -3) indicate introgression between P2 and P3. Hagi: *H. agilis*, Halb: *H. albibarbis*, Hlar: *H. lar*, Hmab: *H. muelleri abbotti*, Hmmu: *H. muelleri muelleri*, Hmol: *H. moloch*, Hpil: *H. pileatus*, Ssyn: *S. syndactylus*.
